# Task rule and choice are reflected by layer-specific processing in rodent auditory cortical microcircuits

**DOI:** 10.1101/860064

**Authors:** Marina M. Zempeltzi, Martin Kisse, Michael G. K. Brunk, Claudia Glemser, Sümeyra Aksit, Katrina E. Deane, Shivam Maurya, Lina Schneider, Frank W. Ohl, Matthias Deliano, Max F. K. Happel

**Author notes:** Correspondence should be addressed to MMZ and MFKH. Author contributions FWO, MD, and MFKH designed research. MMZ, MGKB, and SA collected experimental data. MMZ, MK, CG, KED, SM, and LS analyzed data. MD and MFKH supervised experiments and data analysis. MD established the setup and stimulus protocols. MMZ and MFKH wrote the manuscript. MMZ, CG, MGKB, KD, FWO, MD, and MFKH discussed and all authors reviewed the manuscript.

## Abstract

The primary auditory cortex (A1) is an essential node in the integrative brain network that encodes the behavioral relevance of acoustic stimuli, predictions, and auditory-guided decision making. Previous studies have revealed task-related information being present at both the single-unit and population activity. However, its realization with respect to the cortical microcircuitry is less well understood. In this study, we used chronic, laminar current source density (CSD) analysis from the A1 of behaving Mongolian gerbils (*Meriones unguiculatus*) in order to characterize layer-specific, spatiotemporal synaptic population activity. Animals were trained to first detect and subsequently to discriminate two pure tone frequencies in consecutive training phases in a Go/NoGo shuttle-box task. We demonstrate that not only sensory but also task- and choice-related information is represented in the mesoscopic neuronal population code distributed across cortical layers. Based on a single-trial analysis using generalized linear-mixed effect models (GLMM), we found infragranular layers to be involved in auditory-guided action initiation during tone detection. Supragranular layers, particularly, are involved in the coding of choice options during tone discrimination. Further, we found that the overall columnar synaptic network activity represents the accuracy of the opted choice. Our study thereby suggests a multiplexed representation of stimulus features in dependence of the task, action selection, and the behavioral options of the animal in preparation of correct choices. The findings expand our understanding of how individual layers contribute to the integrative circuit of the A1 in order to code task-relevant information and guide sensory-based decision making.

## Introduction

A central function of the sensory neocortex is the integration of sensory stimulus features and cognitive aspects in behavioral contexts. However, the underlying integrative circuit mechanisms are still only partially understood. In the case of the auditory system, ample evidence has revealed that the primary auditory cortex (A1) integrates sensory information with other contextual and motor signals and further reflects higher cognitive demands responsible for prediction (Kumar et al., 2011; Parras et al., 2017; Town et al., 2018), choice accuracy (Niwa et al., 2012; Caras & Sanes, 2017), and auditory-guided decision making (Brosch et al., 2005; King et al., 2018; Ohl and Scheich, 2005; Tsunada et al., 2015). The salience of behaviorally relevant sounds further critically depends on the exact reinforcement regimes and task rules (Bagur et al., 2018; David et al., 2012; Huang et al., 2019), which renders the auditory cortex a multifarious integrative circuit. These and other studies have described corresponding neural correlates on the level of single neuron or population activity recordings. Some studies have suggested layer-specific differences in the representation of auditory information along the vertical axis of the auditory cortex (Bandyopadhyay et al., 2010; Li et al., 2014; Tischbirek et al., 2019). In agreement, a growing body of human imaging studies has shown attention and task-related modulation of auditory processing in the A1 (Deike et al., 2015; Häkkinen and Rinne, 2018; Petkov et al., 2004; Puschmann et al., 2017) based on gross neural or metabolic response measures. However, how the canonical principles of the columnar processing are reflected in the aforementioned multiplexed function of the A1 is very much unknown (Ohl, 2015; King et al., 2018). We lack a deeper understanding of the underlying neuronal observables on a mesoscopic level characterizing the contributions of whole populations of synaptic circuits across the cortical layers.

Here, we utilized chronic laminar local field potential (LFP) recordings and the analysis of the corresponding current source density (CSD) distribution in the A1 in behaving Mongolian gerbils. Chronic CSD profiles allow the measurement of the spatiotemporal synaptic population activity and to characterize the mesoscopic cortical column physiology. In a first phase animals were trained to detect pure tones of two different frequencies and respond with a ‘Go’ response to both of them (‘Go’ contingency). After successful task acquisition, in a second phase the task rules were changed so that only one of the frequencies remained to be a ‘Go’ signal, while for the other frequency the ‘NoGo’ response was then the correct choice (‘NoGo’ contingency). We found that cortical layers differentially contribute to represent the physical attributes of task-relevant stimuli, the task rule, conditioned motor initiation, behavioral decision making, and the choice accuracy. Based on single-trial CSD data, we used generalized linear-mixed effect models (GLMM) with a logistic link function in order to effectively predict an animal’s behavior. During the initial detection task of two pure tone frequencies all cortical layers contributed to initiate an active avoidancebehavior. The rather task-irrelevant sound frequency was not differentially reflected on a columnar response level. After switching to the more demanding discrimination task employing the same pure tone stimuli, synaptic circuits within mainly granular input layers and supragranular layers reflected the behaviorally observed discriminability between the stimulus classes ‘Go’ and ‘NoGo’. Hence, the task structure affected the columnar representation of auditory information to otherwise identical pure tones. Further, the columnar population activity differed strongly between correct hit responses and correct rejections, while it failed to predict the animal’s behavior during inadequate choices. Our study revealed that the relative contribution of cortical layers to the canonical columnar response is modulated by task-dependent features such as the behavioral relevance of the stimulus, its particular contingency and required action, as well as direct decision variables and the choice accuracy. Finally, this multiplexed information coded by the layer-specific cortical population activity emphasizes the integrative circuit function of the A1 as an instructive mediator between bottom-up routed task-relevant sound features and top-down-controlled auditory-guided decision making.

## Materials & Methods

Experiments were carried out with adult male Mongolian gerbils (*Meriones unguiculatus*, 4 to 8 months of age, 70-90 g body weight, total n=9). All experiments presented in this study were conducted in accordance with ethical animal research standards defined by the German Law and approved by an ethics committee of the State of Saxony-Anhalt.

### Surgery and chronic implantation under electrophysiological control

For chronical *in vivo* electrophysiological recordings a multichannel electrode (Neuronexus, A1×32-6 mm-50-177_H32_21mm) was surgically implanted into the A1. Gerbils were initially anesthetized by an intraperitoneal (i.p.) injection (0.004 ml/g) consisting of 45% ketamine (50 mg/ml, Ratiopharm GmbH), 5% xylazine (Rompun 2%, Bayer Vital GmbH) and 50% of isotonic sodium-chloride solution (154 mmol/1, B. Braun AG). Anesthesia during the surgery was maintained with around 0.15 ml/g*h ketamine i.p. infusion. Anesthetic status was regularly checked (10-15 min) by the paw withdrawal-reflex and breathing frequency. Body temperature was continuously measured and kept stable at 34°C. The primary field A1 of the right auditory cortex was exposed by a small trepanation through the temporal bone (Ø 1mm). This avoids tissue damage and guarantees stable fixation of the implanted electrode on the skull. Another small hole for an initial reference wire (stainless steel, Ø 200-230 µm) was drilled into the parietal bone on the contralateral side. Animals were head-fixed with a screw-nut glued to the rostral part of the exposed nasal bone plate by UV-curing glue (Plurabond ONE-SE and Plurafill flow, Pluradent) that was temporally attached to a metal bar. The recording electrode with a flexible bundle between shaft and connector was inserted perpendicular to the cortical surface into A1 via the small hole.

During the implantation animals were placed in a Faraday-shielded acoustic soundproof chamber. Sounds were presented from a loudspeaker (Tannoy arena satellite KI-8710-32) in 1 m distance to the animal. For verification of the implantation site in A1, a series of pure-tones covering a range of at least 7 octaves were presented (0.25–32 kHz; tone duration 200 ms, inter-stimulus-interval (ISI) 800 ms, 50 pseudorandomized repetitions, sound level 65 dB SPL). Stimuli were generated in Matlab (MathWorks, R2006b), converted into an analog signal by a data acquisition card (sampling frequency 1 kHz, NI PCI-BNC2110, National Instruments), rooted through an attenuator (gPAHGuger, Technologies), and amplified (Thomas Tech Amp75). A measurement microphone and conditioning amplifier were used to calibrate acoustic stimuli (G.R.A.S. 26AM and B&K Nexus 2690-A, Bruel&Kjaer, Germany).

Tone-evoked LFPs were recorded with the multichannel array, pre-amplified 500-fold and band-pass filtered (0.7-300 Hz) with a PBX2 preamplifier (Plexon Inc.). Data were then digitized at a sampling frequency of 1 kHz with the Multichannel Acquisition Processor (Plexon Inc.). Recordings of tone-evoked responses were taken around 30 minutes after implantation and before the final fixation of the electrode. After 30 minutes of laminar recordings, to allow for signal stabilization and verification of the tonotopic location, the electrode and connector (H32-omnetics) were glued onto the animal’s skull with UV-glue. Before enclosing the exposed A1 with UV-glue an antiseptic lubricant (KY-Jelly, Reckitt Benckiser-UK) was applied to the exposed cortex. After the surgery, the wounds were treated with the local antiseptic tyrothricin powder (Tyrosur, Engelhard Arzneimittel GmbH & Co.KG). Directly after the surgery and over the next 2 days, animals received analgesic treatment with Metacam (i.p. 2mg/kg bw; Boehringer Ingelheim GmbH) substituted by 5% glucose solution (0.2 ml). Animals were allowed to recover for at least 3 days before the first session of awake electrophysiological recording.

### Characterization of the recording location in A1 during awake - passive listening

After the recovery period, animals were placed in a 1-compartment box in an electrically shielded and sound-proof chamber in order to re-characterize the tuning properties of the chronically implanted electrode. Acoustic stimuli were presented in a pseudo-randomized order of pure-tone frequencies covering a range of 7 octaves (0.25-16kHz; tone duration: 200 ms, ISI 800 ms, 50 pseudorandomized repetitions, sound level 70 dB SPL), while laminar LFP signals were recorded. Based on the measurements, we observed a rather flat frequency tuning (Suppl. Figure 2B). After each training phase (detection and discrimination) the frequency response tuning was recorded again.

### Shuttle-box training and behavioral paradigm Behavioral paradigm

Operant conditioning was trained in a two-way avoidance shuttle-box task (see Figure 1). The shuttle-box (E15, Coulbourn Instruments) was placed in an acoustically and electrically shielded chamber and contained two compartments separated by a hurdle (3 cm height). We trained animals (n=9) twice a day with a break of at least 5 hours in between both training sessions. In each training session subjects were allowed to habituate for 3 minutes within the shuttle-box. In the first training phase two pure tones with frequencies 1 kHz and 4 kHz were presented both as ‘Go’ conditioned stimuli (CS+). Subjects needed to detect any tone event and respond with a compartment change in order to avoid a mild foot shock (200-500 µA) presented as the unconditioned stimulus (US). We therefore call this phase the detection phase. Within each trial (12-15s), the CS+ tones were repeatedly presented (tone duration 200 ms, ISI of 1.5 s, 70 dB SPL) in a 6 s observation window during which subjects are required to change the compartment in order to make a correct ‘hit’ response.

**Figure 1:**
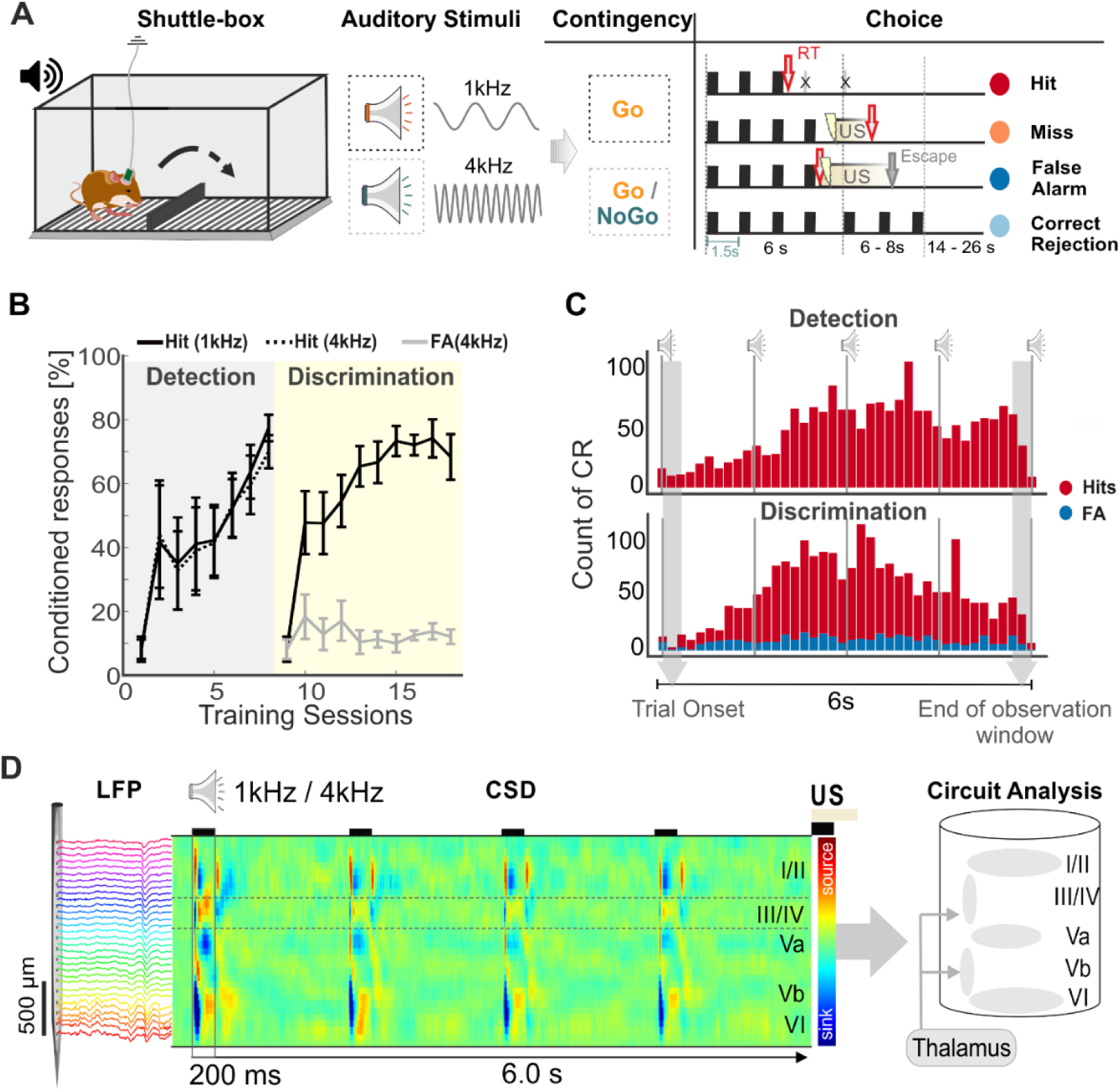
Experimental design, learning curves and chronic CSD recording during auditory-based decision making in a shuttle-box. **A**. Illustration of the two-way avoidance shuttle-box training with chronic recordings in behaving Mongolian gerbils. Subjects were trained to respond to two different pure tone frequencies (1 kHz and 4 kHz; conditioned stimulus - CS) in a Go/NoGo task design to avoid an unconditioned stimulus (US - mild foot shock). During the discrimination phase the contingency of the CS can be either ‘Go’ (CS+) or ‘NoGo’ (CS-) leading to four possible behavioral outcomes (hit, miss, correct rejection – Corr. Rej., false alarm - FA). Right, Illustration of consecutive CS within a trial, length of the observation window (6 s), inter-stimulus interval (1.5 s) and behavioral choices. **B**. Averaged conditioned responses to both CS in the detection and discrimination phase as a function of training sessions. During detection (grey area), hit rates reach almost 80% for both ‘Go’-stimuli (1kHz and 4kHz). At the beginning of the discrimination phase (yellow area), conditioned responses dropped for both stimuli (<10% hit rate). The performance gradually increased reaching again almost 80% for the hit rates and significantly stayed around 20% for the false alarm rates. **C**. Histogram with distributions of the averaged CR reaction times over all trials separately for the detection (top) and discrimination (bottom) phase and hits (red) and false alarms (blue). The majority of CR’s happen after the second CS presentation. **D**. In vivo multichannel LFP recordings were obtained by single-shank silicon probes chronically implanted perpendicular to the surface of the auditory cortex targeting all cortical layers (I – VI). From laminar LFP signals single-trial current source density (CSD) distributions were calculated (here shown is a CSD averaged over 30 repetitions). During CS-presentation (200 ms) tone-evoked CSD components appeared as current sink (in blue) and source (in red) activity reflecting the well-known feedforward information flow of sensory information in the A1 (Happel et al, 2010; 2014)

When subjects shuttled into the other compartment in response to the CS before US onset, this was counted as conditioned response (CR). In case animals did not show a CR within the 6 s observation window this defined a so-called miss trial. Here, the animal received an overlapping presentation of the CS+ and the US until an escape to the other compartment terminated the US/CS presentation. Subjects thereby learned to escape the aversive foot shock within a couple of trials (cf. Happel et al., 2015). In each session we presented each CS+ for 30 times in a pseudorandomized order. The time point at which we changed the task rule from detection to discrimination was oriented at the behavioral performance of each subject individually. For the detection task, which consists only of ‘Go’ trials, the overall task performance of each animal was derived by the d’ values for each session based on signal detection theory. We calculated the d’ as the differences of the z-transforms of the hit rate and the z-transform of the relative inter-trial (ITS) derived from the inverses of a standardized normal distribution function (cf. Happel et al., 2015):

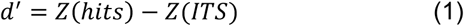

Once animals reached a stable performance of d’> 1 (criterion threshold) for 3 consecutive training sessions, we introduced a change of the task rule and switched to a discrimination task by assigning the former 4 kHz ‘Go’ tone with a ‘NoGo’ (CS-) contingency (n=8; one subject excluded due to epileptic seizure during training). Subjects needed to report on the ‘NoGo’ condition by staying within the compartment to avoid an US, which we call a ‘correct rejection’ (Corr. Rej.). In ‘NoGo’ trials, animals had to stay in the compartment for 12-15 s, while the CS-was continuously played with an ISI of 1.5 s to prevent animals from developing a time estimate of the observation window length over the long training period. If subjects incorrectly crossed within this 12-15 s, the behavioral choice was counted as ‘false alarm’ (FA).

### Acoustic stimuli (CS)

Stimuli were generated in Matlab (MathWorks, R2012b), converted into an analog signal by a data acquisition card (NI PCI-6733, National Instruments), rooted through an attenuator (gPAH Guger, Technologies), and amplified (Black Cube Linear, Lehman). Two electrostatic loud speakers positioned 5cm at both sides of the shuttle-box. A measurement microphone and conditioning amplifier were used to calibrate acoustic stimuli (G.R.A.S. 26AM and B&K Nexus 2690-A, Bruel&Kjaer, Germany).

### Unconditioned stimuli (US)

The mild foot-shock (US) was conditionally delivered by a grid floor and generated by a stimulus generator (STG-2008, Multi-Channel Systems MCS GmbH). Depending on the individual animal sensitivity and performance the shock intensity was adjusted (starting at 200 µA) in steps of 50µA until the escape latencies were below 2s, in order to achieve a successful association of conditioned stimuli (CS) and US (cf. Happel et al., 2015).

### Data analysis

We record all compartment changes during the habituation phase and the training phase. Reaction times of CR’s, escape latencies and the number of inter-trial shuttles (ITS) were recorded. The choice outcomes were characterized as hit, miss, false alarm and correct rejection depending on the animals’ behavior and the contingency of the stimulus (see Figure 1A, right). To evaluate the training progress, we calculate the averaged conditioned response (CR) rates as a function of sessions (Figure 1B).

### Multichannel recordings during training

Multichannel recordings were performed with connecting the head-connector of the animal to a preamplifier (20-fold gain, band-pass filtered, HST/32V-G20; Plexon Inc.) and a data acquisition system (Neural Data Acquisition System Recorder Recorder/64; Plexon Inc.). The cable harness was wrapped by a metal mesh for bite protection. Tension of the cable was relieved by a spring and a turnable, motorized commutator (Plexon Inc.) that permits free movement and rotation of the animal in the box. Broadband signals were recorded continuously using a preamplifier (Plexon REC/64 Amplifier; 1Hz-6 kHz) during the training with a sampling frequency of 12 kHz. Local field potentials were sampled with 2 kHz, visualized online (NeuroExplorer, Plexon Inc. Recording Controller) and stored offline for further analysis. To avoid ground loops between recording system, shuttle-box and the animal we ensure proper grounding of the animal via its common ground and leave the grid floor on floating voltage.

### Analysis of electrophysiological data

#### Current source density (CSD) analysis

Based on the recorded laminar local field potentials, the second spatial derivative was calculated yielding an estimate of the current-source density distribution, as seen in equation:

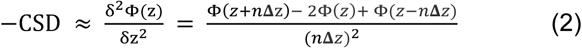

where Φ is the field potential, z is the spatial coordinate perpendicular to the cortical laminae, **Δ***z* is the spatial sampling interval, and n is the differential grid (Mitzdorf, 1985). LFP profiles were smoothed with a weighted average (Hamming window) of 9 channels which corresponds to a spatial kernel filter of 400 µm (Happel et al., 2010).

CSD distributions reflect the local spatiotemporal current flow of positive ions from extracellular to intracellular space evoked by synaptic populations in laminar neuronal structures. CSD activity thereby reveals the spatiotemporal sequence of neural activation across cortical layers as ensembles of synaptic population activity (Mitzdorf, 1985; Happel et al., 2010). One advantage of the CSD transformation that it is reference-free and hence less affected by far-field potentials and referencing artifacts. It allows to observe the local synaptic current flow with high spatial and temporal precision (Kajikawa and Schroeder, 2011). Current sinks thereby correspond to the activity of excitatory synaptic populations, while current sources mainly reflect balancing return currents. The CSD thus provides a functional readout of the cortical microcircuitry function, encompassing a wider, mesoscopic field of view than for instance single-or multi-unit approaches (Buzsáki et al., 2012). Early current sinks in the auditory cortex are therefore indicative of thalamic input in granular layers III/IV and infragranular layers Vb/VI (Happel et al., 2010; Szymanski et al., 2009). In order to describe the overall columnar processing, the CSD profiles were transformed by averaging the rectified waveforms of each channel:

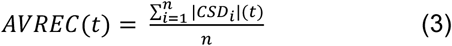

where *n* is the number of recording channels and *t* is time. The AVREC reflects the temporal overall local current flow of the columnar activity (Givre et al., 1994; Schroeder et al., 1998).

### Data Preprocessing

Single-trial data were analyzed via a custom-written graphical user interface (MathWorks, R2016a & R2017b) that visualized the LFP, CSD and behavioral parameters to inspect and mark two types of artifacts: 1) affected recording channels and 2) foot-shock or movement induced signal clipping and distortions. Affected channels were substituted by a linear interpolation method across neighboring, unaffected channels on the level of the LFP (Happel et al., 2010). Shock induced clipping was rejected from the overall signals. Trials with artifacts due to extreme movements were also discarded from further analysis.

### Extraction of signal parameters

Cortical layers were assigned to the recording channels based on the averaged auditory-evoked spatiotemporal CSD current flow in response to the first CS presented during a session and compared to the awake measurement before the training (Suppl. Figure 1). Early dominant current sinks in the auditory cortex are indicative of thalamic input in granular layers III/IV and infragranular layers Vb/VI (Happel et al., 2010; Szymanski et al., 2009) and allow to identify supragranular layers I/II and infragranular layers Va and VI in the CSD recordings (Figure 1D). We determined trial-by-trial root-mean-square (RMS) values of averaged CSD traces within each of the five cortical depths from tone onset of each CS presentation in a time window of 500 ms. Also, we calculated the RMS value of the AVREC within the same time windows for the corresponding overall columnar response. We did not inspect the time-points after a CR, as the CS presentation was terminated. For statistical analysis, single-trial values were z-normalized across trials.

## Statistical Methods

### Statistical test of variance

Statistical difference between groups was tested by one-factorial repeated measures ANOVA (rmANOVA) to account for the hierarchical structure of the data using R Studio (R 3.5.1.). We used an overall significance level of *α* = 0.05 and paired-sample t-tests with a Holm-adjusted significance level (Holm, 1979) for post-hoc testing. Before testing, data was generally z-normalized within each animal and session. The generalized eta squared ƞ^2^_gen_ is reported as measure of effect size calculated using the R package DescTools (Bakeman, 2005; Olejnik and Algina, 2003)

### Mixed-effects logistic regression

For statistical comparison between two-choice classes, parameters of interest were analyzed on a single-trial level using generalized linear-mixed effect models (GLMM) with a logistic link function (cf. Chang et al., 2018) GLMM calculation in in R Studio (R 3.5.1) was done with the lme4 package for model estimation and ggplot2 and sjplot for plotting.

Logistic regression was used for predicting the probability of the binary (0/1) dependent variables ***π***_*i*_ = *E*(***y***_***i***_). The predictions were then wrapped by the logistic link function:

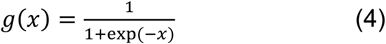

to map the predictions of the model to the interval between 0 and 1. In the mixed-effects logistic regression, random effects were additionally introduced to model subject-specific variance by:

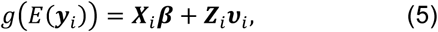

where ***y***_*i*_ is the vector of all responses of the *i*^*th*^ animal, ***X***_***i***_ and ***Z***_***i***_ are design matrices, ***β*** the fixed effects and ***v***_***i***_ the animal-specific random effects. The parameters of the estimated model can be interpreted as logarithmic odds ratios 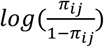, where *π*_*ij*_ corresponds to the probability of the outcome to be 1 for animal *i* in trial *j*. The GLMM thus allows for an intuitive interpretation of its predicted values (choice probabilities) and its estimated coefficients (logarithmic odds ratios). As such, GLMMs are optimally suited to compare data on a trial-by-trial-level while accounting for within-subject variability. Random intercepts were introduced to account for the general variability in overall activity across subjects and random slopes to allow for the fixed effect to vary between animals. We z-normalized the AVREC RMS values for the GLMM to facilitate the estimation procedure.

### Evaluation of the model

Calculation of the marginal (R^2^m) and conditional (R^2^c) coefficient of determination was done using the MuMIn package (Barton, 2019). The R^2^m represents the variance in the dependent behavioral variable explained by the fixed effect of the respective CSD variable (across subjects), while the R^2^c reflects the total variance explained by the model’s fixed and random effects, respectively (Muff et al., 2016). In a binary GLMM, the R^2^m is independent of sample size and dimensionless, which allows comparing fits across different datasets (Nakagawa and Schielzeth, 2013).

## Results

### Auditory decision making with change of the task rule in a shuttle-box

We trained Mongolian gerbils in an auditory cued two-way active avoidance shuttle-box task to respond to two pure tones (of frequencies 1 and 4 kHz) presented as conditioned stimuli, while recording local field potentials from the primary auditory cortex using laminar multichannel electrodes (Figure 1). Gerbils were trained in two separate phases. First, in a detection training phase, both CS were assigned with a ‘Go’ contingency and required subjects to change the compartment to actively avoid the unconditioned stimulus (mild electric foot shock). We trained animals over consecutive sessions until they reached a stable detection of both stimuli significantly above chance level (Figure 1A,B). Thereby, we yielded sufficient data for both behavioral choices. In the subsequent training phase, the contingency of the 4 kHz pure tone was changed to a ‘NoGo’ stimulus (CS-), while 1 kHz was maintained as CS+ (or ‘Go’ stimulus). During this phase, animals needed to discriminate the two pure tones in order to avoid the US. Here, we classified behavioral choices depending on the response of the animal and the contigency as hit, miss, correct rejection, or false alarm (Figure 1A,B). Averaged conditioned response curves across training sessions for both training phases showed significant improvement of task performance (Figure 1B). During detection training averaged hit rates reach almost 80% for both ‘Go’-stimuli (1 kHz and 4 kHz). During the initial discrimination phase, conditioned response rates dropped significantly for both stimuli (<10% hit rate). Hence, animals did not transfer the behavioral choice for the 1 kHz pure tone from the detection phase, but completely abandoned their detection-based avoidance strategy. They quickly re-associated the 1 kHz CS with a ‘Go’ contingency and showed increasing hit responses within 1-2 sessions, while false alarm rates in response to the ‘NoGo’ 4 kHz tone were significantly lower (∼10-20%). Over the entire training procedure, the reaction times were found to be mainly after the first CS presented within a trial. Compartment changes started to increase in response to the second CS and were equally distributed over the subsequent 4.5 seconds of the observation window (Figure 1C). This suggests that the task design allows the subjects to use at least the presentation of a second CS to evaluate their planned behavioral choice.

### Task rule impacts on the columnar representation of sound frequency

Over the entire training, multichannel LFP recordings were obtained by single-shank silicon probes chronically implanted in the primary auditory cortex (Figure 1**Error! Reference source not found**.D). In an averaged CSD trace, the tone-evoked activity in response to the repetitive CS presentation, reflecting the spatiotemporal feedforward flow of sensory information across cortical layers in the A1 (Happel et al., 2010; 2014), marked the most prominent laminar response pattern. During detection, we generally observed highly similar CSD patterns in response to the two pure tones (both ‘Go’ stimuli) with respect to the spatiotemporal current flow (Figure 2A). Initial current sink activity was observed within granular layers III/IV and infragranular layer Vb reflecting cortical depths of main thalamocortical inputs from the ventral medial geniculate body. Subsequent synaptic activity is then routed to supragraular layers I/II and infragranular layers Va and VI. The overall columnar response exceeded the 200 ms duration of the pure tone presentation. In awake passive listening subjects CSD profiles were generally also very similar in response to both pure tones (Suppl. Figure 2A), which is due to considerably similar and flat frequency tuning properties across the entire group of animals measured (Suppl. Figure 2B).

**Figure 2:**
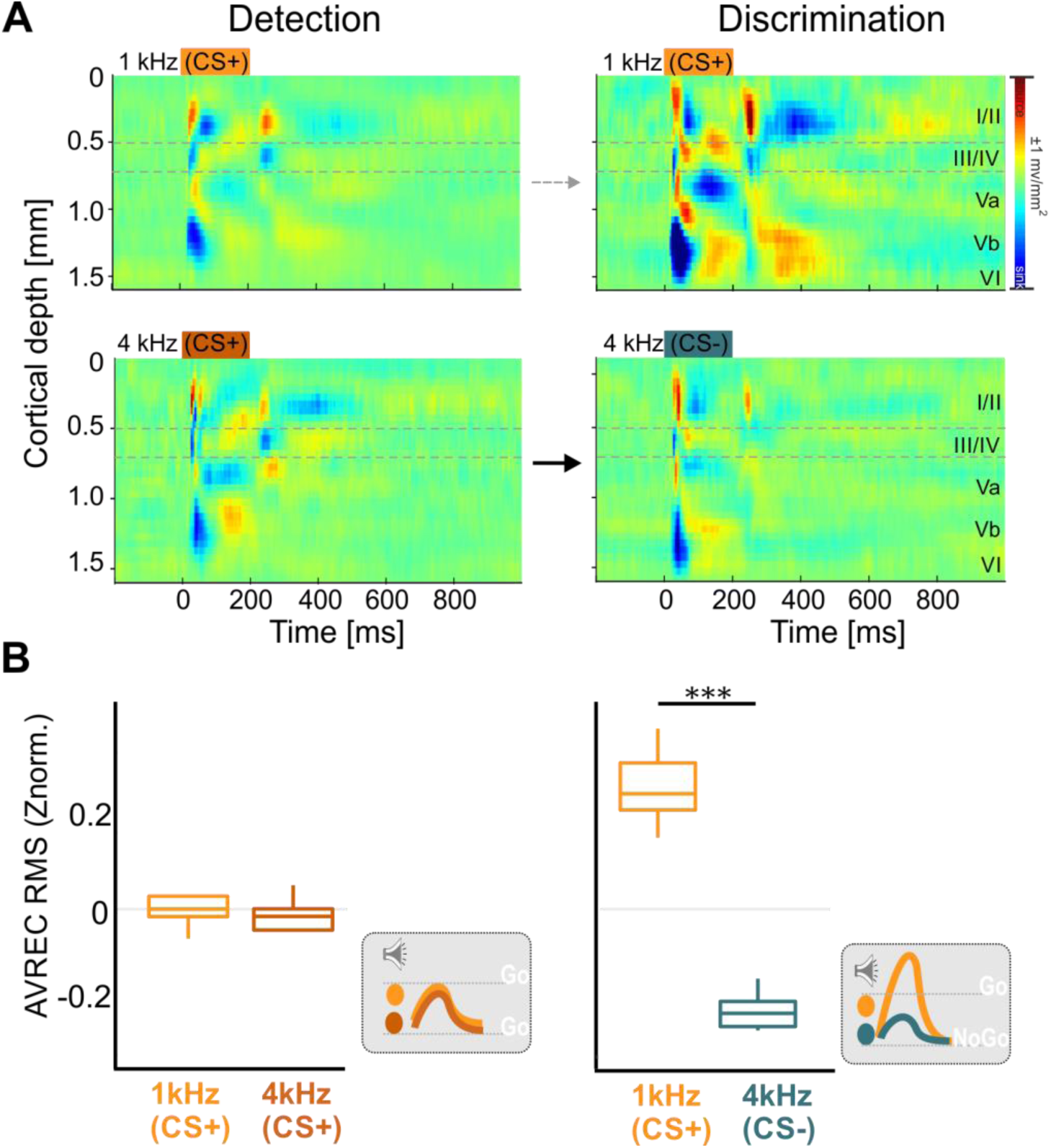
Stimulus-related activity during different training phases. **A**. Representative example of an averaged CSD profile across all trials of the detection (*left*) and discrimination (*right*) phase of one subject. The CSD profiles show the tone-evoked activity after the first presentation of both conditioned stimuli within a trial (*top*: 1 kHz, *bottom*: 4 kHz; tone duration: 200 ms; indicated by dashed bar in upper left panel). Evoked CSD patterns between the two pure tones frequencies showed no obvious differences during the detection phase but yielded considerably different CSD patterns during discrimination for the CS+. **B**. RMS values of the AVREC (time window of 500 ms beginning at each tone presentation and z-normalized) shown for each of the four consecutive CS and separated by the different behavioral outcomes during the two task phases. Box plots represent median (bar) and interquartile range, and bars represent full range of data. Significant bar indicate differences revealed by pairwise testing. Schematic illustration of the evoked cortical activity in dependence of stimulus frequency and task rule are shown in grey inserts.

During the discrimination phase, however, the two physically identical stimuli evoked considerably different CSD patterns. While the overall tone-evoked columnar activity in Go-trials showed a marked increase, the activity in NoGo-trials was rather unchanged or slightly decreased (Figure 2). In order to quantify the overall columnar activity strength, we compared the root mean square values of the AVREC (AVREC RMS; z-normalized) calculated for the entire trace in each trial (Figure 2B). A one-way repeated-measures ANOVA (rmANOVA) revealed that during the detection phase the overall activity over the trial between the two CS+ did not differ (*F*_*1,8*_ = 0.20, *p* = 0.668). During discrimination, the CS+ evoked significantly more cortical overall current flow compared to the CS- (*F*_*1,7*_ = 143.63, *p* < 0.001). Accordingly, our findings show that the activation strength of the auditory cortex in response to pure tones depends on the task rule (Figure 2B, grey insets).

### Auditory cortex represents behavioral choice and choice accuracy

We further differentiated how the cortical recruitment depends on the decision taken by the animal. We compared AVREC RMS values during a 500 ms window beginning with each CS onset. At each of the CS+ presentation during detection training, one-way rmANOVAs consistently report a significant relation of the z-normalized AVREC RMS values to hits and misses at 1 kHz and 4 kHz (1^st^ CS+: *F*_3,24_ = 5.71, *p* = 0.004, 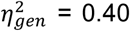; 2^nd^ CS+: *F*_3,24_ = 38.15, *p* < 0.001, 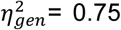; 3^rd^ CS+: *F*_3,24_ = 34.36, *p* < 0.001, 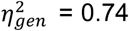; 4^th^ CS+: *F*_3,24_ = 28.02, *p* < 0.001, 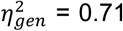; for details see Suppl. Table 1). Holm-corrected post-hoc pairwise t-tests revealed that the evoked AVREC RMS value (z-norm.) after the first CS presentation is similar for hit and miss trials, with only a small difference between hits and misses at 4 kHz. Consecutive CS+ presentations evoked significantly higher RMS values during hit trials compared to miss trials (Figure 3, *left*). These findings were independent of the actual stimulation frequency (1 or 4 kHz). In the discrimination phase, we found a differential recruitment of auditory cortex columnar activity depending on frequency and choice of the subjects at all CS presentations (1^st^ CS+: *F*_3,21_ = 8.75, *p* < 0.001, 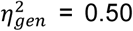; 2^nd^ CS+: *F*_3,21_ = 29.44, *p* < 0.001, 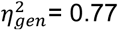; 3^rd^ CS+: *F*_3,21_ = 25.32, *p* < 0.001, 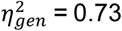; 4^th^ CS+: *F*_3,21_ = 34.94, *p* < 0.001, 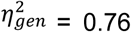; Suppl. Table 1) (Figure 3A, *right*). Cortical activation in the 500 ms time window around the first CS showed only minor differences in the post-hoc tests and was significantly lower during correct rejection trials compared to hit and false alarm trials. During later CS presentations throughout the trial, we found a stable pattern of columnar activity. During hit trials, cortical activation was significantly highest compared to all other classes. Correct rejections showed the lowest cortical recruitment. In contrast, cortical activation during miss and false alarm trial did not differ at any CS presentation throughout the trial. Note that cortical recruitment was generally stronger during trials in which animals reported a compartment change for both CS individually, comparable to findings in the detection phase. However, as the cortical activity during incorrect miss trials and false alarm trials did not differ significantly, the variability of cortical activation in our data seemed to not be explained by a mere correlate of motor responses or motor planning, but also depended on the contingency of a stimulus. Indeed, the strongest difference observed was between the two correct choice options of the animal, namely between hits and correct rejections. Hence, cortical recruitment during detection was influenced to a larger degree by the behavioral action taken by the animal, rather than the physical stimulus characteristic of tone frequency. During discrimination, cortical recruitment was influenced by both, the frequency, coding for the contingency of the stimulus, and the choice accuracy of the taken action (Figure 3B). For a further substantiation of this finding, see the following paragraph.

**Figure 3:**
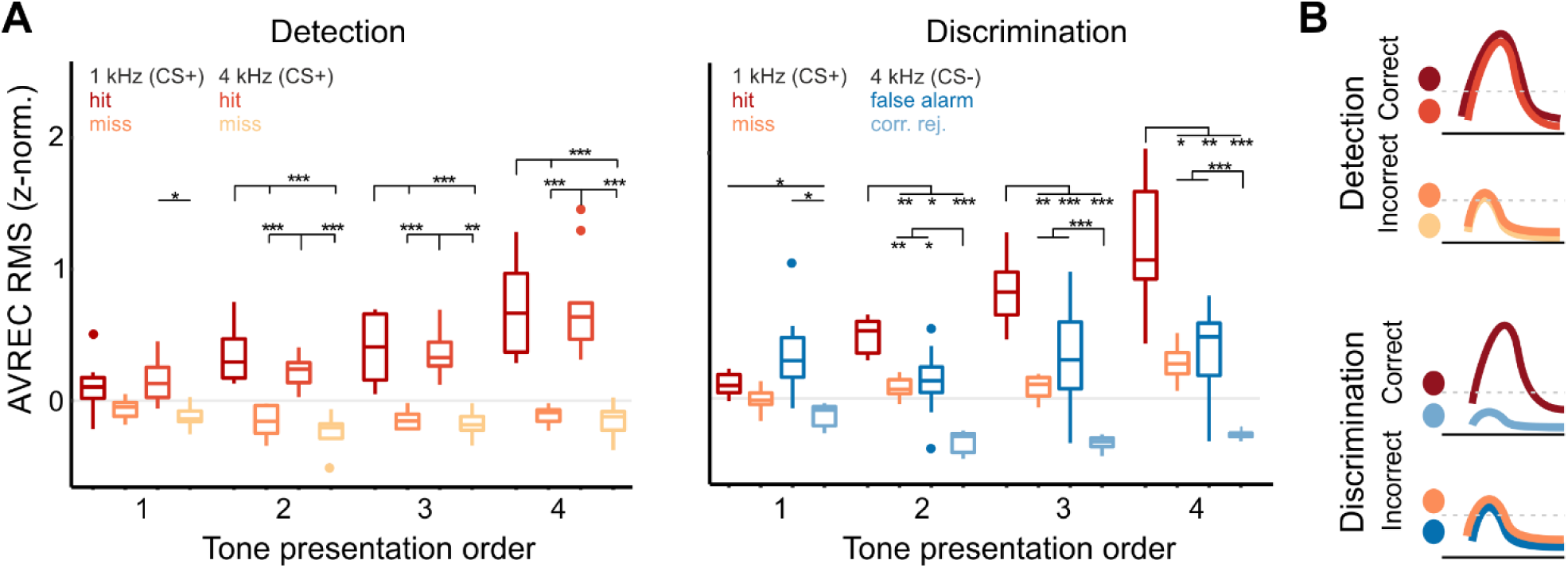
Behavioral choices and contingency are both reflected in population activity of the A1. Averaged AVREC RMS values (500 ms window at CS onsets) plotted with respect to the conditioned stimuli and behavioral choice. **A**. *Left*, During the detection phase evoked activity was significantly higher during hit trials compared to miss trials independent of the stimulation frequency. *Right*, In the discrimination phase, cortical activity was strongest during correct hit trials and lowest during correct rejections. During trials of incorrect behavioral choices (miss/false alarm) tone-evoked activity was characterized by intermediate amplitudes and did not differ. Box plots represent median (bar) and interquartile range, and bars represent full range of data. Dots represent outliers. Significant bars indicate differences revealed by a one-way rmANOVA (see Suppl. Table 1) **B**. In summary, cortical activity was generally higher in trials in which animals showed a conditioned response in comparison to trials where animals stayed in the compartment. Cortical activity differed strongest between correct behavioral choices, namely hits and correct rejections.

### Representation of contingency is layer-specific and differs with task rule

In order to investigate the contribution of cortical layers to the observed effects, we analyzed binary classes on a single-trial level using generalized linear-mixed effect (GLMM; Chang et al., 2018). The GLMM analysis revealed that in the detection phase, the AVREC trace RMS (z-norm.) was not dependent on the presented frequency of the two conditioned stimuli (*left, R*^*2*^*m* = 0, *ns*.; Figure 4, *left*). During the discrimination phase, an increase in the AVREC trace RMS was a reliable predictor that the 1kHz ‘Go’ stimulus was presented (*R*^*2*^*m* = 0.17, *p* < 0.001; Figure 4, *right*). Hence, the columnar activity in auditory cortex in response to the same conditioned stimuli differed in dependence of the task and was only separable when both had contrasting contingencies. We further applied the GLMM to the RMS value measured over the entire trace activity within single cortical layers (I/II, III/IV, Va, Vb, VI) in order to reveal the source of the aforementioned results on a layer-specific level (Figure 4). In the detection phase, the two CS+ used as binary class in the GLMM could not predicted significantly for any particular cortical layer. When we applied the GLMM for the two conditioned stimuli during the discrimination phase, now reflecting the distinct contingency of CS+ and CS-, we observed a moderate prediction of the model with an R^2^m of 0.12 for only the granular input layers. Detailed results for each model are reported in Suppl. Table 2.

**Figure 4:**
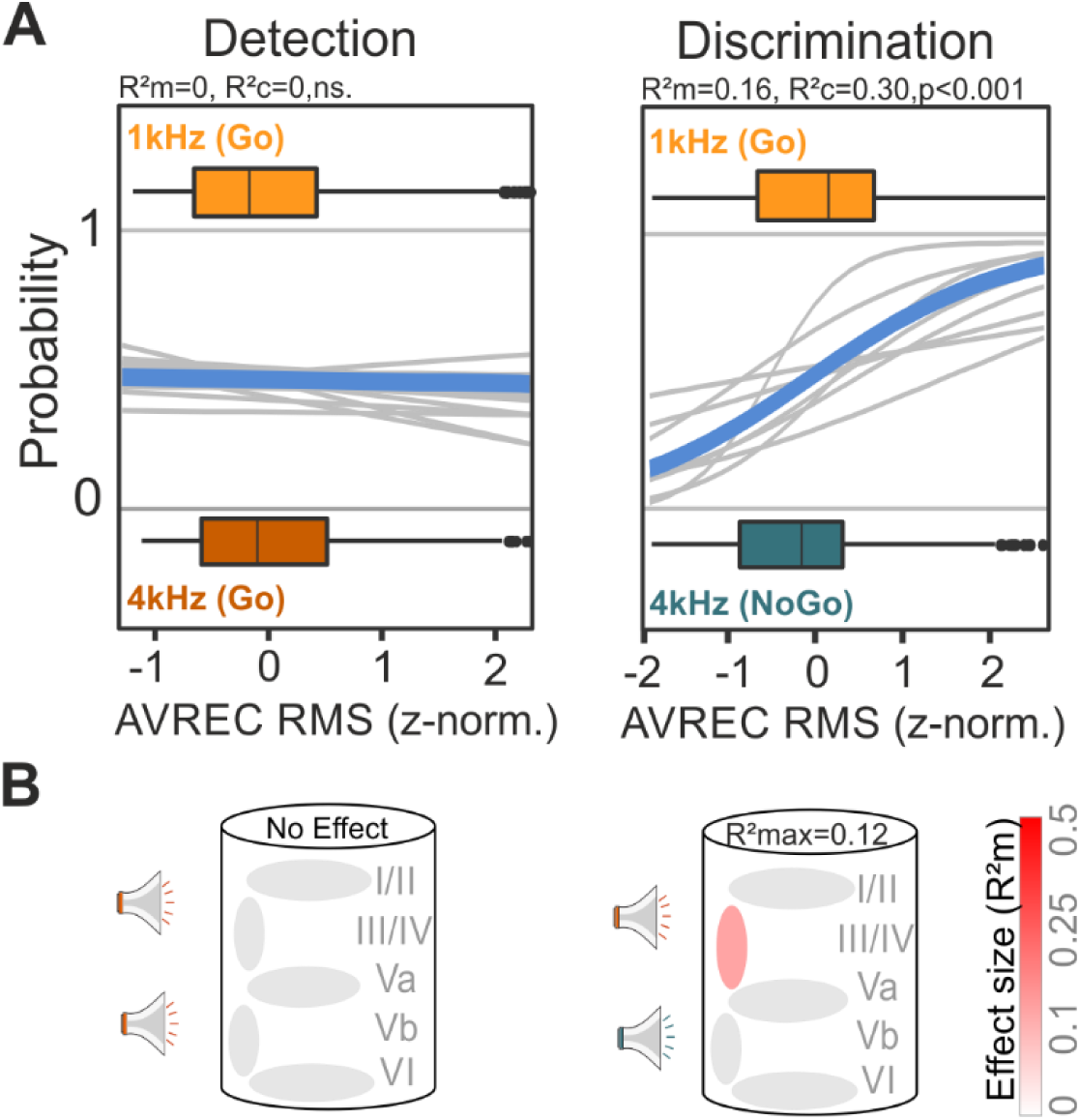
Representation of contingency, not frequency revealed in synaptic population activity of granular input layers. Parameters of interest were analyzed on a single-trial level using generalized linear-mixed effect models **A**. Logistic regression curves show the probabilities of the presented CS (1kHz and 4kHz as the dependent variable) for individual subjects (gray) and as an average (blue). The boxplots above and below the curves represent the mean (bar), interquartile range (box) the full range of data (whiskers). The AVREC trace RMS did not predict the frequency of the conditioned stimuli (1 kHz and 4 kHz) during the detection phase (*left*, R^2^m=0, R^2^c=0, ns.). During discrimination an increase in the AVREC trace RMS significantly indicated that the 1 kHz ‘GO’ stimulus was played (R^2^m=0.16, R^2^c=0.30, p<0.001). Hence, auditory cortical activity in response to the same conditioned stimuli differed in dependence of the task. **B**. GLMM’s were applied to RMS values measured within single cortical layers (I/II, III/IV, Va, Vb, VI). The illustration of the cortical column below indicates the GLMM predictability based on data from corresponding layers to the binary behavioral choice combinations. The color illustrates the effect size for the model-based R^2^m (grey= no effect to red=strong effect). The top R^2^m value (R^2^max) depicts the best fit result for all layers tested. In the detection phase, the two CS+ used as binary class in the GLMM revealed no significant prediction for any particular cortical layer. During the discrimination phase we observed a moderate prediction of the model with R^2^m = 0.12 for the granular input layers. The detailed results for each GLMM are reported in Suppl. Table 2.

In a next step, we used the GLMM to predict the behavioral choices rather than the stimulus frequency (Figure 5). Therefore, we compared the AVREC RMS values (z-norm.) of the 500 ms windows around the tone presentation that preceded an active avoidance response of the animal (hit/false alarm) or around the last CS in the observation window in trials without a CR (miss/cor. rej.). During the detection phase, a higher AVREC RMS was a robust predictor for trials with a correct hit response compared to miss trials with lower overall cortical activity (*R*^*2*^*m* = 0.32, *p* < 0.001; Figure 5A, *left*). In order to test the contribution of distinct cortical layers to the coding of different behavioral choices, GLMM predictions were calculated for RMS values of each layer separately. We found that during the detection phase, cortical activity in infragranular layers were good predictors (*R*^*2*^*m* = 0.25 – 0.35, p < 0.001), while supragranular and granular layer were less accurate (*R*^*2*^*m* = 0.11 – 0.19, p < 0.001; Figure 5B, *left*; cf. Suppl. Table 3).

**Figure 5:**
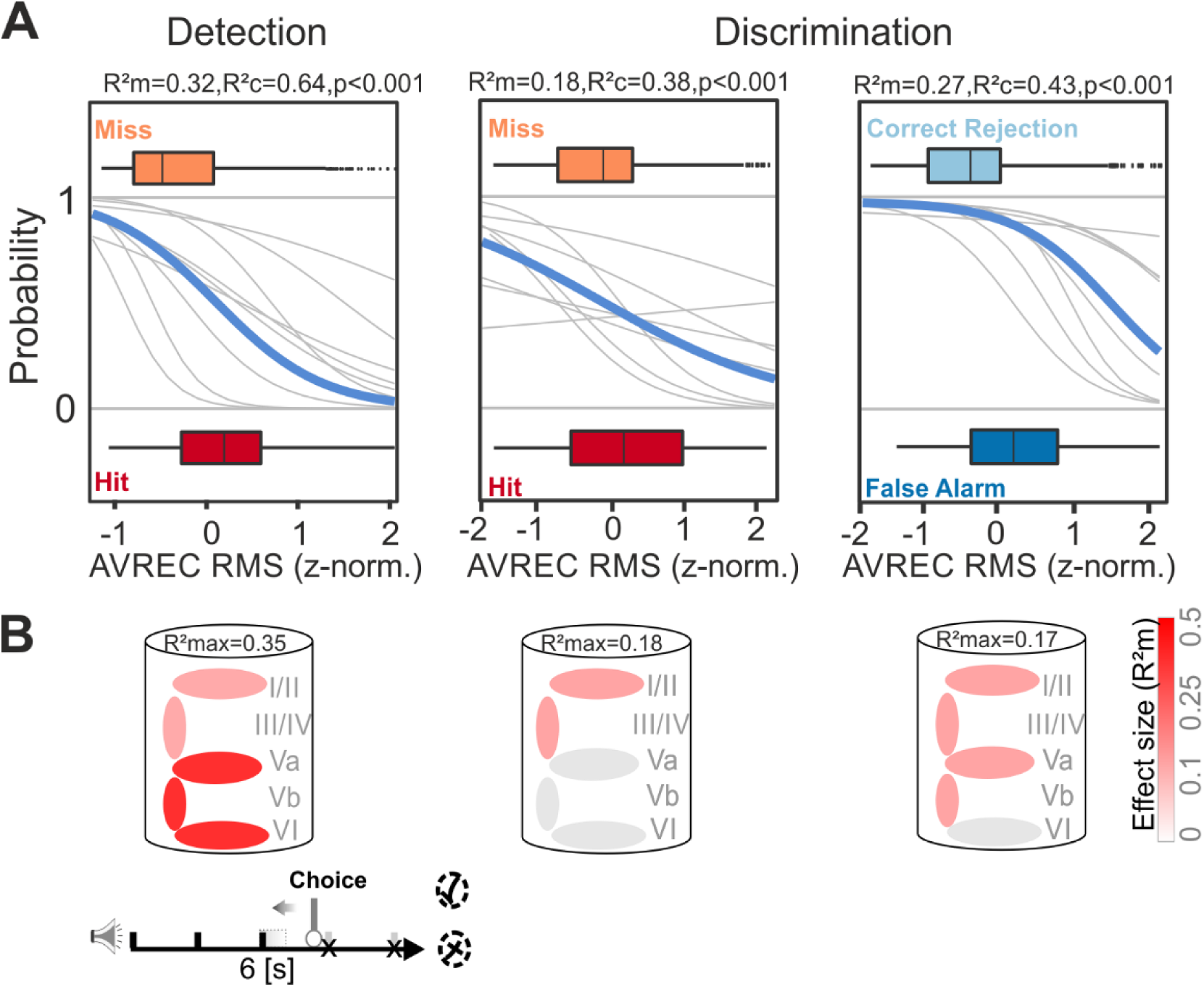
Layer-specific contribution to behavioral choice. **A**. GLMM and logistic regression analysis was used to predict the behavioral choice of the subjects. *Left*, During the detection phase RMS values of the AVREC (z-norm.) in the 500 ms time window around the CS presentation which was initiating a hit response was significantly higher compared to the fourth CS during miss trials (R^2^m=0.32, p<0.001). *Middle*, this was also true for the discrimination phase, although with only a moderate effect size (R^2^m=0.18, p<0.001). When comparing data from ‘NoGo’ trials, false alarm and correct rejections could be predicted with a high effect size (R^2^m=0.27; p<0.001). **B**. GLMM predictions for each layer showed that cortical activity from all layers were good predictors (R^2^m=0.1-0.35), especially higher effect we observed at deeper layers Va, Vb, and VI, for the two possible choices (hit/miss). This finding was independent of the actual spectral content of the presented stimulus (1 kHz/4 kHz; see Figure 3). During the discrimination phase, granular and supragranular layers appear to be important for the differential representation of the behavioral choice in ‘Go’-trials (R^2^m=0.14-0.18). For ‘NoGo’-trials, the GLMM revealed that false alarms are accompanied by significantly higher activity in all cortical layers except of layer VI compared to correct rejections (R^2^m=0.17, p<0.001). Supragranular layers were also the best predictor between false alarms and correct rejections classes. The detailed results for each GLMM are reported in Suppl. Table 3.

During the discrimination phase the AVREC RMS was also predicting the choice outcome during ‘Go’-trials (hits vs. misses) with a moderate effect size (*R*^*2*^*m* = 0.18, R^2^c=0.38, p < 0.001; Figure 5A, *right*). For ‘NoGo’-trials the GLMM was able to predict the outcome with a high effect size: false alarms were effectively predicted by stronger cortical recruitment than measured during correct rejections (*R*^*2*^*m* = 0.27, *p* < 0.001; Figure 5A, *right*). During discrimination, granular and supragranular layers appear to be important for the differential representation of the behavioral choice in ‘Go’-trials (R^2^m = 0.14-0.18). During ‘NoGo’-trials, the RMS value of all cortical layers except of layer VI were good predictors for the trial outcome (*R*^*2*^*m* =0.10-0.17, *p*<0.001), while supragranular layers were also the best predictor between false alarms and correct rejections (see Suppl. Table 3).

### Choice accuracy is robustly represented throughout cortical layers

In a last step we compared correct and incorrect choice options of the subjects showing that only correct choices lead to a distinguishable cortical circuit activation (Figure 6). The AVREC RMS could predict the outcome in correct trials with a high effect size: Correct ‘hit’ responses can be predicted by higher RMS values of the AVREC trace in the time window before the actual decision compared to the time window at the trial end during correct rejections (R^2^m=0.45, p<0.001). In contrast, the two incorrect choices ‘false alarms’ and ‘miss’ were not predictable by the GLMM (R^2^m=0.04; n.s.; Figure 6A). The layer-specific analysis further revealed that particularly supragranular layer activity contributed to the differential cortical activation between the correct choice classes (R^2^m=0.18-0.51; p<0.001; cf. Suppl. Table 4; Figure 6, *left*). Nevertheless, all cortical layers were recruited in a distinct way, so that the whole cortical column differs in activity during correct hit and correct rejection choices. In accordance with the insignificant GLMM result on the overall columnar activity measured by the AVREC RMS, also no cortical layer activity could predict the two incorrect choices (false alarm/miss; Figure 6, *right*).

**Figure 6:**
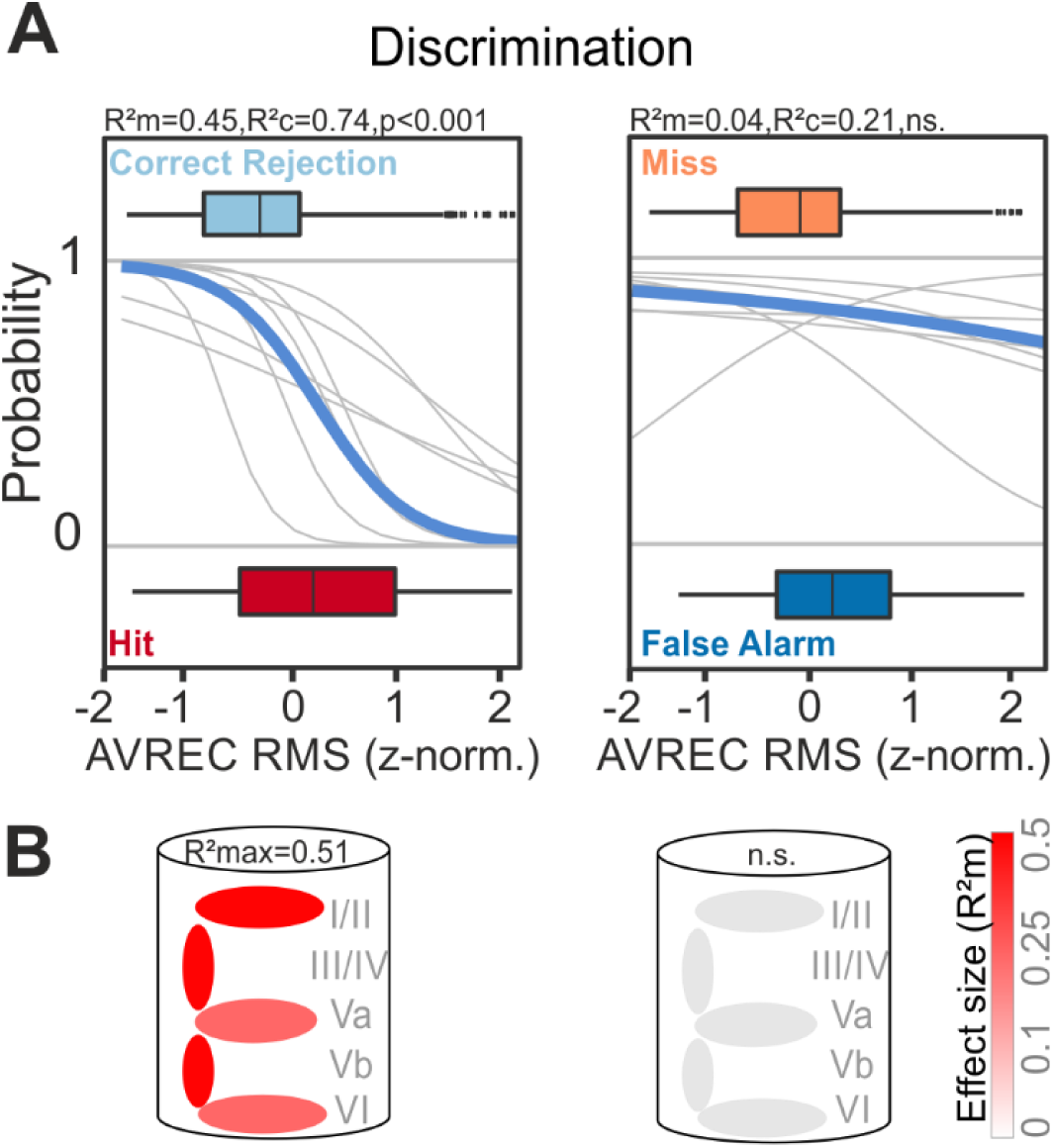
Representation of choice accuracy across layer-specific population activity in A1. **A**. Predictability of correct (*left*) and incorrect (*right*) choices during the discrimination phase were modelled by GLMM and logistic regression. Correct ‘hit’ responses can be predicted by higher RMS values of the AVREC trace in the time window before the actual decision compared to the time window at the trial end during correct rejection responses (R^2^m=0.45, p<0.001). In contrast, the two incorrect choices ‘false alarms’ and ‘miss’ were not predictable by the GLMM (R^2^m=0.04; n.s.). **B**. Activity from all cortical layers contributed to the differential cortical activation between the correct choice classes, while the largest effect size was found for supragranular layers (R^2^m=0.51; p<0.001). In accordance with the insignificant GLMM result on the overall columnar activity measured by the AVREC, also no cortical layer activity could predict the two incorrect choices (false alarm/miss). The detailed results for each GLMM are reported in Suppl.Table 4.

## Discussion

In this study, we chronically recorded local field potentials and calculated current source density (CSD) distributions from the primary auditory cortex of Mongolian gerbils. Animals were trained to first detect and then to discriminate two pure tone frequencies in two consecutive training phases in a Go/NoGo shuttle-box task. Based on the laminar distribution of CSDs, we demonstrate that not only sensory but also task- and choice-related information is represented in the neuronal population activity distributed across cortical layers. The frequencies of two pure tones used as conditioned stimuli were only differentially represented in the A1 when they differed in their contingency, i.e. when their discrimination was behaviorally relevant for the task. Cortical activity also differed with action selection generally showing a higher recruitment during trials where the animal initiated a compartment change. During the detection phase, infragranular layers contributed most to those differences. In contrast, recruitment of synaptic activity in supragranular layers was the most robust predictor for choice outcomes during the discrimination phase (Figure 5). We further found a robust representation of the choice accuracy independent of the actual action selection of the animal. While all cortical layers showed a stronger recruitment during correct hit trials compared to correct rejections, we did not observe differences between false alarms and misses (Figure 6). Hence, our findings argue for a multiplexed representation of stimulus- and task-related features distributed across the cortical layers.

### Supragranular layer activity better classifies the stimulus contingency than the presented tone frequency

We use auditory instrumental conditioning with detection and consecutive discrimination of two pure tone frequencies centered within the optimal hearing range of the gerbil. After successful detection, animals had to abandon their initially learned strategy and ‘re-associate’ one of the two CS with a new meaning during the discrimination phase. We found that such a switch of the task rule caused the animals to completely abandon the previous but still valuable ‘knowledge’ about parts of the stimulus representation and to re-learn a new set of behavioral action-outcome contingencies (Figure 1B).

We found that during discrimination stimulus-dependent features between Go- and NoGo-stimuli are represented differentially, as the sound frequency gathered a behavioral relevance due to the shift of the task rule. A GLMM analysis revealed that the representation of two different pure tone frequencies is distinguishable on the level of the A1 population activity only if there is the behavioral need to discriminate both stimuli. These task-dependent representations emerge as accumulating evidence throughout the trial and are most strongly represented right before a behavioral choice of the animal (see Figure 3). During the discrimination phase, animals needed to differentially represent the sound frequency of the two conditioned stimuli to successfully perform the task. Henceforth, during this training phase, the need of spectral integration was likely to be behaviorally more important. Here, we found that particularly input layers III/IV and supragranular layers I/II were more strongly recruited during trials that led to an active conditioned response. Activity during hits and false alarms was higher compared to misses and correct rejections, respectively. This might reflect the need for more crosscolumnar communication within supragranular layers in order to integrate the spectral content of a presented CS necessary to promote the correct behavioral choice (Hickmott & Merzenich, 1998; Sakai & Suga, 2002; Happel et al., 2014; Francis et al., 2018).

Therefore, we propose that the representation of stimulus features in sensory cortex, such as tone frequency in A1, does not depend alone on the transmission process of the sensory information via the primary sensory pathways, but is significantly modulated by the behavioral need and the behavioral relevance of a stimulus. Such influence is based presumably on higher order top-down inputs from, for instance, parietal and frontal areas (Caras and Sanes, 2017; Polley et al., 2006; Rodgers and DeWeese, 2014; Runyan et al., 2017; Steinmetz et al., 2019).

### Correlates of motor initiation dominate A1 population activity during detection

Motor initiation has been reported before to enhance or suppress sensory-driven activity in other (primary) sensory cortices depending on region, system and task-engagement (Busse et al., 2017; Steinmetz et al., 2019). From our data we hypothesize, that, during the detection, the tone-evoked activity in the primary auditory cortex may be modulated by auditory-guided motor initiation (Brosch et al., 2015; Huang et al., 2019; Niwa et al., 2012a). The distinct sound frequency of a pure tone seems less determining on the activity strength. Deep output layer activity (layers Va-VI) showed a significant increase of activity during hit trials. This is in accordance with the findings that neurons in these layers convey information to downstream motor centers, as the basal ganglia or the striatum, which play an important role for the control of motor decisions by the sensory cortex (Xiong et al., 2015; Znamenskiy and Zador, 2013). Further, the selection of an appropriate action might also be conveyed directly to motor cortex via direct anatomical projections (Matyas et al., 2010; Huang et al., 2019). Ample evidence argues that our findings reflect a motor-related modulation of the cortical physiology, rather than a movement artifact. In our data, auditory cortex activity reflected the initiation of motor actions during detection learning most prominently in deeper layers. Hence, motor-related signals were reflected on a layer-specific level while showing a conserved spatiotemporal profile of the tone-evoked CSD, which is in strong favor of a motor-related modulation of the cortical physiology. A muscle correlate, as a far field artifact, would have affected all recording channels. We controlled this by a trial-by-trial analysis excluding such trials (cf. Figure 2A). Here, the reference-free CSD measurement might be an effective filter. During discrimination, the cortical activity was less accurate in predicting motor response initiation but was more accurate during correct choice options (Figure 5 and 7). Cortical population activity did not differ during false alarm and miss trials. However, cortical activity was elevated between the consecutively presented CS during hit trials. This argues for an accumulative evidence about the stimulus contingency that the animals kept persistently over the trial, which was instructive for an auditory-guided action. These differences between hit and false alarms argue that the motor-related preparatory signal cannot fully explain the variability in our data set. Rather, we find a combinatorial representation of stimulus contingency, task rule, selection accuracy, and motor initiation that accumulates in its richness over the experimental procedure of the actual decision.

### Choice accuracy is represented throughout the cortical column

The modulation of the cortical activity by contingency and motor initiation reflects a cortical correlate of choice accuracy: in the discriminant Go/NoGo-paradigm, we found all cortical layers to be more strongly activated during correct hits compared to correct rejections (Figure 6). In contrast, cortical activity during false alarms and misses did not differ. Hence, the cortical representation of spectral information during discrimination training (see Figure 3) is further dependent on the accuracy of the promoted behavior. While several studies also observed an enhanced representation of target stimuli that initiated an auditory-guided motor response in various Go/NoGo discrimination tasks (Bagur et al., 2018; Fritz et al., 2003; Gold et al., 1999), others found higher cortical recruitment during correct rejections compared to hit trials in a Go/NoGo task in the macaque A1 (Huang et al., 2019). Previous findings demonstrate the potentially inherent neuronal variability comprised of the exact task design at hand, aversive or appetitive reinforcing regimes, and stimulus characteristics which may partially explain contradictory findings (David et al., 2012; Osmanski and Wang, 2015) and needs further evaluation. Another relevant aspect is the temporal relation of the observed effects to the repetitive tone presentation throughout the trial in our task design. We focused our analysis on time windows of 500 ms around the consecutively presented CS which covered the sensory-dominated columnar response (Figure 2A) and preceded the behavioral choice. Other reports of choice-related activity in the auditory cortex during discrimination of tone events also reported that such representation accumulates until the animal’s decision (Bizley et al., 2013; Niwa et al., 2012). We further analyzed a time window of 500-1000 ms after each stimulus presentation in order to separate the relative modulation of cortical layer activity by sensory-driven effects from the task-related, but potentially temporally distributed information (data not shown). This analysis revealed that the choice accuracy is represented across all cortical layers as accumulating evidence across the entire trial length (cf. Bizley et al., 2013) and hence is present also independently from the stimulus-dominated auditory response.

Altogether, our study demonstrates that the auditory cortex population activity reflects the task complexity at hand, as well as the choice accuracy of the animal. Motor initiation has a stronger impact on cortical activity during detection training, where other task-dependent features, such as coding of the contingency, are absent. During the more complex discrimination task, other factors also affect cortical activity. Overall, our results show that the layer-specific population activity in the sensory cortex is highly dependent on the behavioral task and accordingly reflects the performance of the individual subject within different task phases.

## Conclusions

Previous work and the current study show that neuronal activity already in the primary auditory cortex encodes sounds in ways that are directly relevant to behavior. We found that the entire ensemble activity of the A1 columnar circuits closely represented task-relevant stimulus features, the task rule and behavioral choice variables suggesting its instructive role for auditory-guided decision making. While infragranular layers dominated the cortical processing modes during action selection within a detection context, supragranular layers gained relevance when stimulus features needed to be integrated during discrimination. Our study thereby expands our understanding of the layer-specific cortical circuit processing modes which code task-relevant information in order to guide sensory-based decision making and behavioral adaptation during strategy change. We have now begun to reveal the functional computations performed by single neurons and of the local and long-range cortical networks they are integrated in (cf. Happel, 2016). Future studies will enunciate the more widespread brain networks for mediating perceptual decision making, in which the A1 circuitry reflects only one important hub.

## Acknowledgements

We would like to thank Kathrin Ohl and Anja Guerke for their technical assistance. This project was founded by the Deutsche Forschungsgemeinschaft (DFG SFB 779) and by the Leibniz Association (LIN Postdoctoral Network, LPN).

## Supplemental Information

**Supplementary Figure 1:**
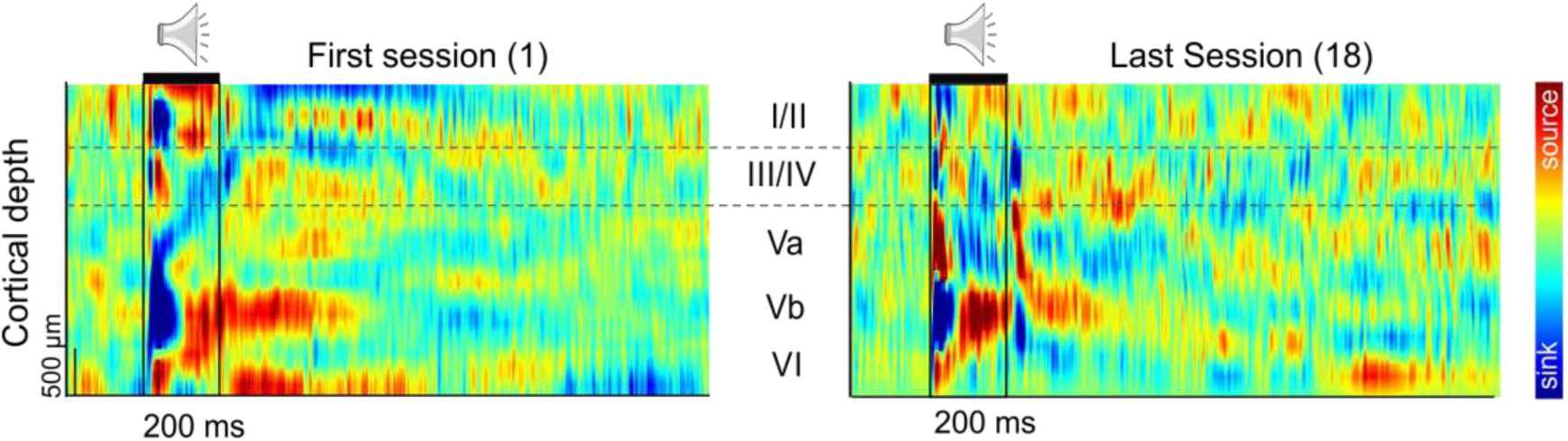
Long-term stability CSD recordings from all cortical layers in A1. Representative example of an averaged CSD profile from one subject of the first training session (detection; *left*) and the last discrimination session (*right*). Based on the averaged auditory-evoked activity in response to the first presentation of the conditioned stimuli within a trial (time window: 1500ms; tone duration: 200 ms; indicated by the black frames) we assign the cortical input layers (I/II – VI) to the respective recording channels (indicated with the dashed lines). The example illustrates the stability of the electrode positioning over the course of the training.

**Supplementary Figure 2:**
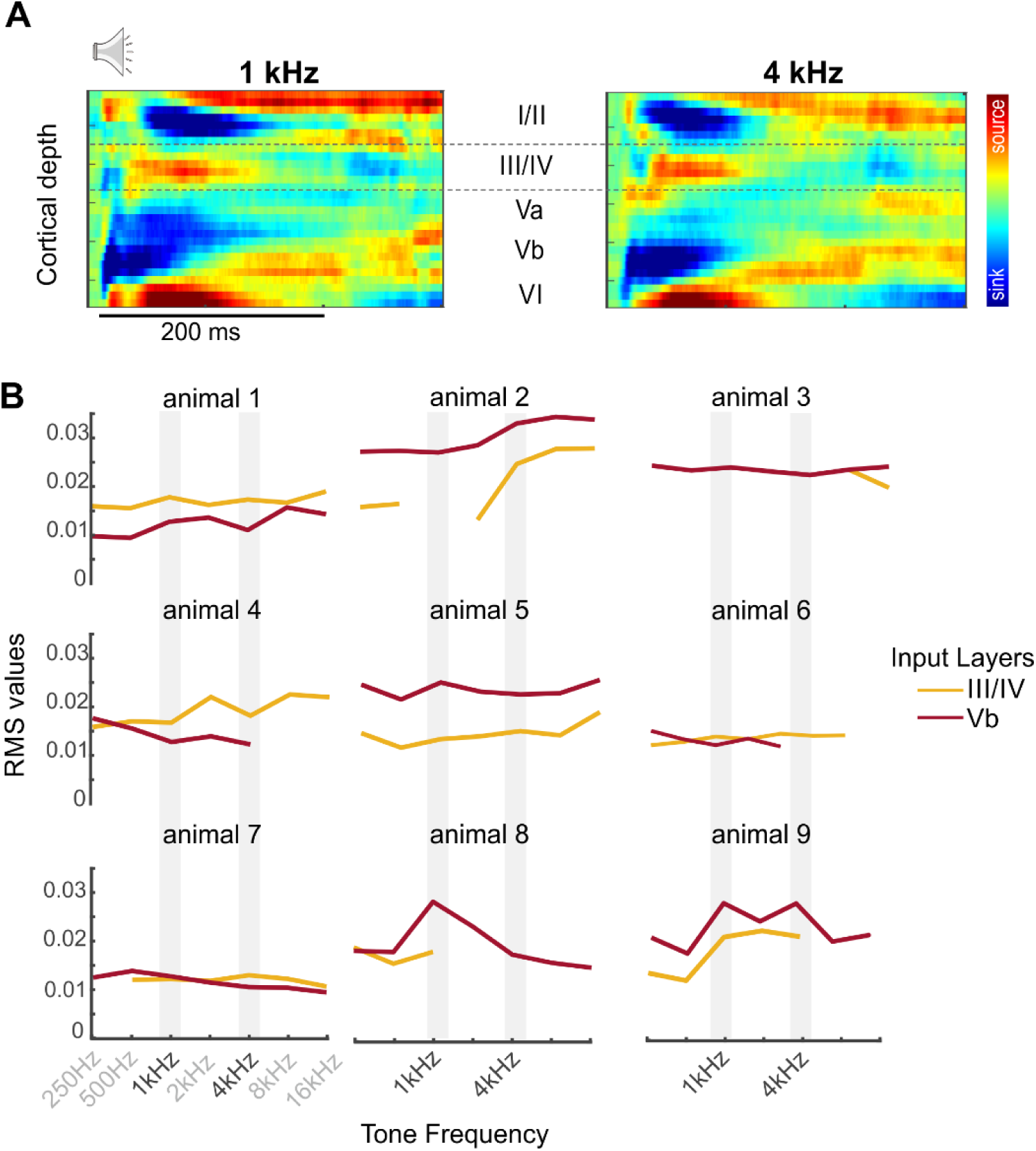
Characterization of tuning properties in the recording location A1. **A**. Representative example of an averaged CSD profile during the first awake, but passively listening measurement before the actual start of the behavioral training (see Methods and Materials; n=1). CSD activity is shown for the two pure tone frequencies also used during the later training, namely 1 kHz, *left* and 4 kHz, *right* (tone duration: 200 ms, ISI 800 ms, 50 pseudorandomized repetitions, sound level 70 dB SPL). **B**. Tuning curves of layer-specific CSD RMS amplitudes receiving early thalamocortical inputs (III/IV and Vb; n=9). Mean of CSD RMS values (averaged over 50 trials per frequency) of cortical layers III/IV and Vb are plotted as a function of stimulation frequency revealing flat frequency tuning in awake, passively listening subject (cf. Deane et. al (2019) *bioRxiv*; doi:https://doi.org/10.1101/810978).

**Suppl. Table 1.** (cf. Figure 3): rmANOVA of choice- related contingencies AVREC RMS.

|  |  |  |  |
| --- | --- | --- | --- |
| **Detection** |  |  |  |
| **1<sup>st</sup> stimulus** | $F_{3,24} = 5.710$ | $p = 0.004$ | $\eta_{gen}^2 = 0.40$ |
| **2<sup>nd</sup> stimulus** | $F_{3,24} = 38.15$ | $p < 0.001$ | $\eta_{gen}^2 = 0.75$ |
| **3<sup>rd</sup> stimulus** | $F_{3,24} = 34.36$ | $p < 0.001$ | $\eta_{gen}^2 = 0.74$ |
| **4<sup>th</sup> stimulus** | $F_{3,24} = 28.02$ | $p < 0.001$ | $\eta_{gen}^2 = 0.71$ |
| **Discrimination** |  |  |  |
| **1<sup>st</sup> stimulus** | $F_{3,21} = 8.750$ | $p < 0.001$ | $\eta_{gen}^2 = 0.50$ |
| **2<sup>nd</sup> stimulus** | $F_{3,21} = 29.44$ | $p < 0.001$ | $\eta_{gen}^2 = 0.77$ |
| **3<sup>rd</sup> stimulus** | $F_{3,21} = 25.32$ | $p < 0.001$ | $\eta_{gen}^2 = 0.73$ |
| **4<sup>th</sup> stimulus** | $F_{3,21} = 34.94$ | $p < 0.001$ | $\eta_{gen}^2 = 0.76$ |

**Suppl. Table 2.**
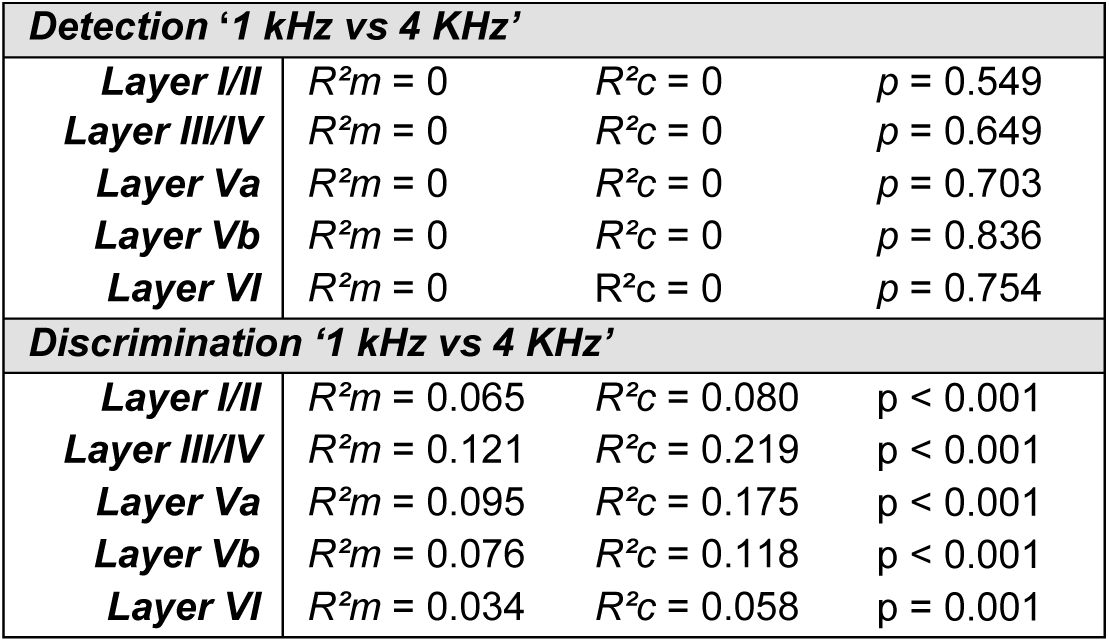
(cf. Figure 4B): GLMM layer specific applied to the conditioned stimuli 1 kHz vs 4 kHz.

|  |  |  |  |
| --- | --- | --- | --- |
| **Detection '1 kHz vs 4 KHz'** |  |  |  |
| **Layer I/II** | $R^2m = 0$ | $R^2c = 0$ | $p = 0.549$ |
| **Layer III/IV** | $R^2m = 0$ | $R^2c = 0$ | $p = 0.649$ |
| **Layer Va** | $R^2m = 0$ | $R^2c = 0$ | $p = 0.703$ |
| **Layer Vb** | $R^2m = 0$ | $R^2c = 0$ | $p = 0.836$ |
| **Layer VI** | $R^2m = 0$ | $R^2c = 0$ | $p = 0.754$ |
| **Discrimination '1 kHz vs 4 KHz'** |  |  |  |
| **Layer I/II** | $R^2m = 0.065$ | $R^2c = 0.080$ | $p < 0.001$ |
| **Layer III/IV** | $R^2m = 0.121$ | $R^2c = 0.219$ | $p < 0.001$ |
| **Layer Va** | $R^2m = 0.095$ | $R^2c = 0.175$ | $p < 0.001$ |
| **Layer Vb** | $R^2m = 0.076$ | $R^2c = 0.118$ | $p < 0.001$ |
| **Layer VI** | $R^2m = 0.034$ | $R^2c = 0.058$ | $p = 0.001$ |

**Suppl. Table 3.** (cf. Figure 5B): GLMM layer-specific applied to the behavioural choices.

|  |  |  |  |
| --- | --- | --- | --- |
| **Detection 'Hit vs Miss'** |  |  |  |
| **Layer I/II** | $R^2m = 0.113$ | $R^2c = 0.537$ | $p = 0.056$ |
| **Layer III/IV** | $R^2m = 0.190$ | $R^2c = 0.406$ | $p < 0.001$ |
| **Layer Va** | $R^2m = 0.351$ | $R^2c = 0.624$ | $p < 0.001$ |
| **Layer Vb** | $R^2m = 0.258$ | $R^2c = 0.569$ | $p < 0.001$ |
| **Layer VI** | $R^2m = 0.200$ | $R^2c = 0.396$ | $p < 0.001$ |
| **Discrimination 'Hit vs Miss'** |  |  |  |
| **Layer I/II** | $R^2m = 0.144$ | $R^2c = 0.240$ | $p < 0.001$ |
| **Layer III/IV** | $R^2m = 0.188$ | $R^2c = 0.361$ | $p = 0.001$ |
| **Layer Va** | $R^2m = 0.007$ | $R^2c = 0.149$ | $p = 0.216$ |
| **Layer Vb** | $R^2m = 0.049$ | $R^2c = 0.152$ | $p = 0.001$ |
| **Layer VI** | $R^2m = 0.033$ | $R^2c = 0.212$ | $p = 0.118$ |
| **Discrimination 'False alarm vs Correct Rejection'** |  |  |  |
| **Layer I/II** | $R^2m = 0.164$ | $R^2c = 0.465$ | $p < 0.001$ |
| **Layer III/IV** | $R^2m = 0.170$ | $R^2c = 0.318$ | $p < 0.001$ |
| **Layer Va** | $R^2m = 0.192$ | $R^2c = 0.480$ | $p = 0.003$ |
| **Layer Vb** | $R^2m = 0.105$ | $R^2c = 0.263$ | $p < 0.001$ |
| **Layer VI** | $R^2m = 0.096$ | $R^2c = 0.239$ | $p < 0.001$ |

**Suppl. Table 4.** (cf. Figure 6B): GLMM layer specific applied to the choice accuracy.

|  |  |  |  |
| --- | --- | --- | --- |
| **Discrimination 'Hit vs Correct rejection'** |  |  |  |
| **Layer I/II** | $R^2m = 0.517$ | $R^2c = 0.607$ | $p < 0.001$ |
| **Layer III/IV** | $R^2m = 0.435$ | $R^2c = 0.573$ | $p < 0.001$ |
| **Layer Va** | $R^2m = 0.181$ | $R^2c = 0.426$ | $p = 0.001$ |
| **Layer Vb** | $R^2m = 0.378$ | $R^2c = 0.479$ | $p < 0.001$ |
| **Layer VI** | $R^2m = 0.193$ | $R^2c = 0.287$ | $p < 0.001$ |
| **Discrimination 'Miss vs False Alarm'** |  |  |  |
| **Layer I/II** | $R^2m = 0.164$ | $R^2c = 0.465$ | $p = 0.104$ |
| **Layer III/IV** | $R^2m = 0.170$ | $R^2c = 0.318$ | $p = 0.004$ |
| **Layer Va** | $R^2m = 0.192$ | $R^2c = 0.480$ | $p = 0.202$ |
| **Layer Vb** | $R^2m = 0.105$ | $R^2c = 0.263$ | $p = 0.058$ |
| **Layer VI** | $R^2m = 0.096$ | $R^2c = 0.239$ | $p = 0.322$ |

